# A multimodal learning approach for automated detection of wildlife trade on social media

**DOI:** 10.1101/2025.09.24.678024

**Authors:** Mohammad Momeny, Ritwik Kulkarni, Jooel Rinne, Andrea Soriano-Redondo, Enrico Di Minin

## Abstract

Social media data and machine learning methods for automated content analysis are increasingly being used in ecology and conservation science. A current limitation is the lack of methods for automated multimodal analysis of textual and visual content among other data modalities. In this study, we introduce a multimodal content analysis method applied to the investigation of wildlife trade on YouTube. Our approach consists of analyzing text through transformer based neural networks and video keyframes using convolutional neural networks as part of multimodal filtering followed by classification where a decision fusion module identifies instances of wildlife trade. The decision fusion module achieved an F-score of 0.72 among textual classifiers for trade detection and of 0.77 among visual classifiers for species identification. This multimodal classification helped detect wildlife trade in 3,715 out of 86,321 filtered YouTube posts, featuring 226 species for sale, including 51 Critically Endangered, 62 Endangered, 60 Vulnerable, 25 Near Threatened, and 28 Least Concern species. The proposed multimodal learning methods can be used more broadly for other ecological and biodiversity conservation applications.

**The bigger picture:** The unsustainable trade in wildlife is a major driver of biodiversity loss, threatening thousands of species across the Tree of Life. While online platforms have become popular spaces for advertising wildlife and exotic pets for sale, monitoring these platforms remains extremely challenging. Traditional surveillance methods are not scalable, and automated tools have typically focused on either text or image analysis in isolation, limiting their effectiveness in identifying nuanced instances of wildlife trade. Our study introduces a multimodal machine learning framework that integrates textual and visual data to detect potential wildlife trade on YouTube. By combining natural language processing with deep learning for image analysis, and filtering millions of posts down to those most relevant, our method significantly improves detection accuracy. This dual-layered approach uncovered thousands of posts featuring hundreds of species, many of which are threatened. This work demonstrates how advances in machine learning can support ecological monitoring and conservation by providing timely, data-driven, insights into online trade networks. In the pursuit of reducing biodiversity loss, this study offers an approach for bridging the gap between online behavior and real-world ecological outcomes.

**Highlights:**

- Introduces a multimodal content analysis approach for detecting wildlife trade on YouTube by integrating textual and visual data.
- A multimodal filtering technique reduces irrelevant text and video content, enhancing analytical efficiency.
- A decision fusion module then combines results from text and video filtering improving wildlife trade detection accuracy.
- The proposed methods are applicable across multiple online platforms and suitable for diverse tasks in ecology and biodiversity conservation.

## Introduction

Earth biodiversity is in steep decline, with abundance and number of wild species decreasing worldwide (Díaz et al., 2019). Unsustainable exploitation of species is one of the main drivers of the biodiversity crisis, as millions of animals and plants, and their derived products, are traded unsustainably every year at a global scale (Haken, 2011). Estimates from the International Union for the Conservation of Nature (IUCN) indicate that 24% of the >31,500 evaluated species of terrestrial birds, mammals, amphibians, and squamate reptiles are currently traded (IUCN, 2001; Scheffers et al., 2019). Recent estimates for vertebrates show that where trade occurs, population numbers can potentially decrease by 62% (Morton et al., 2021). Besides vertebrate species, wildlife trade affects thousands of less knownlesser-known species across the Tree of Life (e.g., plants, fungi, and invertebrates (Fukushima et al., 2020). Unsustainable wildlife trade can also impact the wider ecosystems, for example, by modifying trophic networks, changing communities, or by inducing the establishment of invasive species (Hughes et al., 2023). Its impacts can even reach human societies, through the transmission of zoonotic diseases, with catastrophic health and economic consequences (Cardoso et al., 2021).

Digital and social media platforms have become a popular venue for selling wildlife (Lavorgna, 2014; Moloney et al., 2023; Salas-Picazo et al., 2023; Sung et al., 2021; Wyatt et al., 2022; Xu et al., 2020). Evidence points to wildlife trade occurring across taxa, platforms, and sites (Marshall et al., 2020; Soriano-Redondo et al., 2023). For example, over 200 reptile, mammal, bird, amphibian, and invertebrate species were advertised for sale on two Facebook profiles in Togo (Harrington et al., 2021) Another study found 599 relevant advertisements originating from 41 websites focusing on 43 endemic and range-restricted reptile species from the Lesser Antilles (Rinne et al.). An analysis of butterflies’ trade on eBay detected 50,555 transactions involving 3,767 species (Wang et al., 2023).

Efficient investigation of online wildlife trade requires using machine learning methods (Di Minin et al., 2019). Thus far, methods have focused on automatically identifying species and wildlife products, which are known to be traded online, from images using computer vision (Cardoso et al., 2023; Di Minin et al., 2018; Kulkarni et al., 2025). More recently, machine vision methods have been adapted to detect images of exotic pets on sale based on the non-natural environments in which they are located (Kulkarni & Di Minin, 2023). Natural language processing methods have also been used to automatically identify online wildlife trade content through the analysis and synthesis of digital text (Kulkarni & Di Minin, 2021). These methods allow for the recognition and extraction of locations, organizations, species, quantities, and prices mentioned in digital posts. They also allow to monitor and study the sentiment for and reactions of online wildlife trade content (Di Minin et al., 2019; Fink et al., 2020). Multimodal methods that can simultaneously identify text and image content are still vastly missing, but they could help improve the automated identification of online wildlife trade content further. Images can sometimes misleadingly imply that an item is for sale – for instance, when animals are pictured in artificial environments like zoos or urban areas (Tøn et al., 2024). Similarly, text analysis alone may fall short in identifying species for sale, as language often contains numerous homonyms that can confuse a species and other items (e.g., *Jaguar* the vehicle brand vs *jaguar* the feline).

Here, we introduce an approach for the multimodal analysis of text and video content, aimed at investigating the online wildlife trade on YouTube. YouTube is being used by wildlife traders and pet owners to showcase and market wildlife species and products (Carvalho et al., 2023; Fink et al., 2021; Haq et al., 2023; Siriwat & Nijman, 2018) as it offers users the ability to upload, share, and view a vast array of video content (Ek & Samahita, 2023). Our approach consists of analyzing text through transformer based neural networks and video keyframes using convolutional neural networks as part of multimodal filtering followed by classification where a decision fusion module identifies instances of wildlife trade (Figure 1). During the preprocessing phase, irrelevant textual and visual data are filtered out using a cascading multimodal filtering approach (video filtering after text filtering) to enhance performance and processing speed. Additional deep neural networks are then applied to analyze the filtered text data and video keyframes. Finally, a decision fusion module that integrates the outputs of both textual and visual models is used to detect instances of wildlife trade. To enable multimodal analysis, we developed a custom YouTube video collection pipeline targeting content that referenced species listed in Appendix I of the Convention on International Trade in Endangered Species of Wild Fauna and Flora (CITES). A subset of the collected dataset was manually annotated into four categories: Explicit Wildlife Trade, Wildlife Trade Information, Other Wildlife Content, and Irrelevant. These annotated samples were then used to train a classifier model and validate filtering algorithms at each stage of the multimodal classification framework, enabling detection of wildlife trade. See the Methods section for a full explanation.

**Figure 1.**
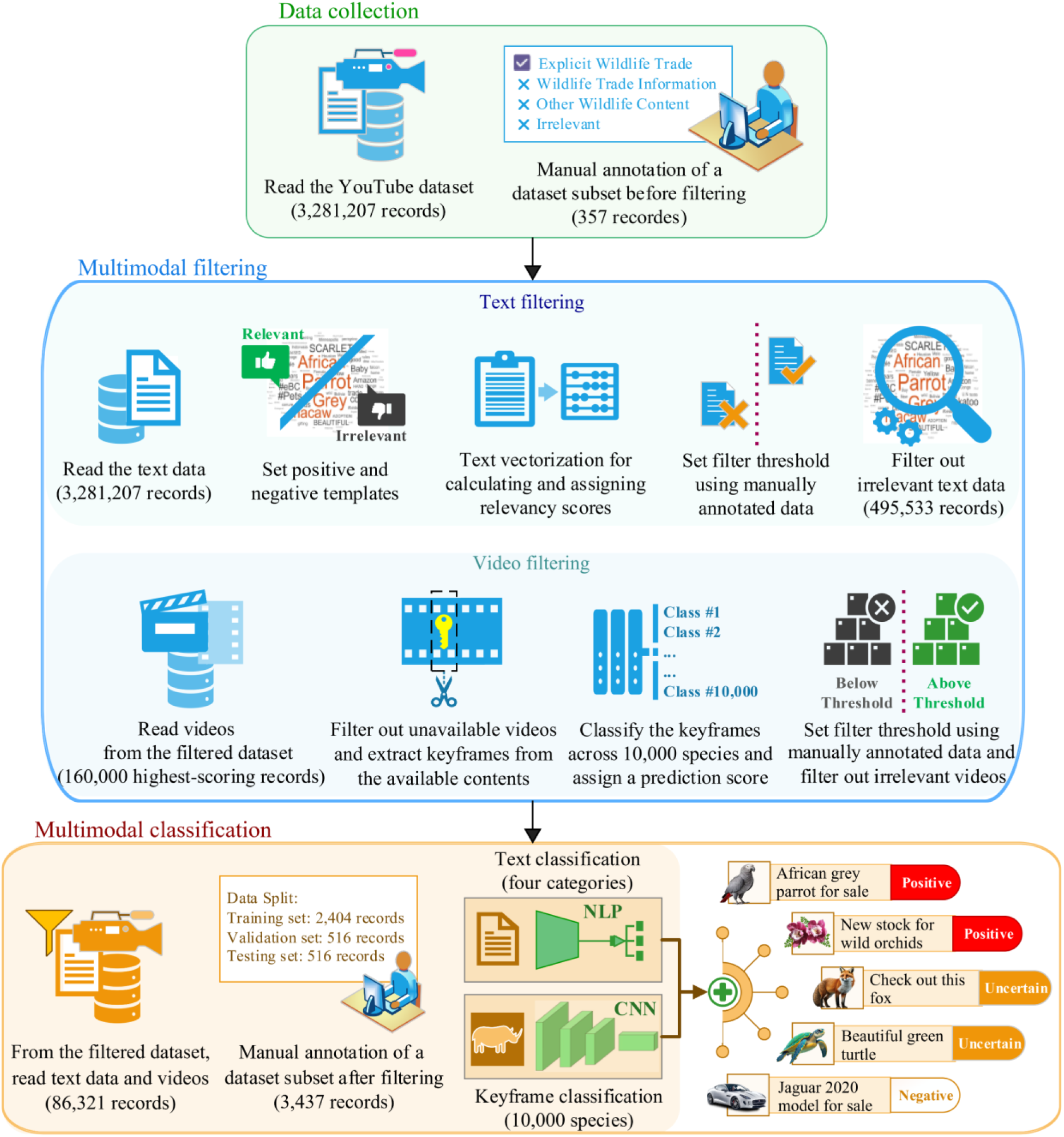
Schematic pipeline of the proposed multimodal learning approach.

## Results

The performance of the proposed multimodal filtering pipeline for detecting relevant wildlife trade content was assessed via a manual annotation of its outputs across textual and visual modalities. Different subsets of the original dataset (N=3,281,207) were generated by sequential application of filtering algorithms and each subset was further subsampled for manual annotation, to calculate the percentages of relevant and irrelevant categories. Initial manual assessment of a random subset of data resulted in “Explicit Wildlife Trade” category comprising 1.12% of the subset. A new subset of data was sampled by applying the text filtering algorithm that assigns a relevancy score to each post and selecting a random set of posts that scored above the relevancy threshold, this improved the percentage of “Explicit Wildlife Trade” to 2.93%. Further, 364 posts were sampled that had the highest relevancy score from text filtering algorithm and manually assessed again. This resulted in 5.79% of the posts to be “Explicit Wildlife Trade”. Lastly, another subset of posts was selected that had top relevancy scores from text filtering and were then filtered using image analysis of the associated videos. A manual assessment found 8.61% of this sample to be “Explicit Wildlife Trade”. Conversely, irrelevant content dropped sharply from 66.29% in the original dataset to 22.61% after relevancy-based text filtering, 6.89% after top score-based text filtering, and just 3.54% after filtering based on image analysis of the associated video with the posts (Table 1). The substantial reduction in irrelevant content demonstrates the effectiveness of the pipeline featuring multimodal filtering approach of first filtering based on text and then on image analysis, in eliminating noise. It is important to note that although the text filtering algorithm is seemingly having lower number of explicit sales post hits (a natural statistics of the data on the web), it is highly successful in removing “non-species” related posts from the data and thereby reducing the computational burden of downstream algorithms like video filtering which can focus on more probable posts for wildlife trade.

**Table 1.** Distribution of YouTube post categories before and after multimodal filtering.

| **Category** | **Original dataset** | **Text filtering** | **Text filtering**<br>**Highest-scoring** | **Video filtering** |
| --- | --- | --- | --- | --- |
| Total Records | 3,281,207 | 495,533 | 160,000 | 86,321 |
| Sample Size (Random Selection) | 356 | 376 | 364 | 393 |
| Explicit Wildlife Trade | 1.12% | 2.93% | 5.79% | 8.61% |
| Wildlife Trade Information | 0.28% | 5.59% | 1.10% | 2.28% |
| Other Wildlife Content | 32.30% | 68.88% | 86.23% | 85.57% |
| Irrelevant | 66.29% | 22.61% | 6.89% | 3.54% |

After the last stage of image-based filtering of videos associated with the posts, 86,321 posts were marked as highly probable for wildlife trade related content. Out of these, 3,437 randomly selected posts were manually annotated for both the text content into four categories (Explicit Wildlife Trade, Wildlife Trade Information, Other Wildlife Content, and Irrelevant) and image content (presence/absence of a species in the video). The manual annotation was used to train neural net algorithms, separately for text and images extracted from the video. For textual classification, we implemented two different methods, All-MPNet-Base-V2 (transformer architecture) and LSTM. All-MPNet-Base-V2 consistently outperformed LSTM in classifying textual features, achieving a sensitivity of 0.69, specificity of 0.98, and F-measure of 0.72 (Table 2). For image analysis, we evaluated DarkNet53, InceptionResNetV2, and EfficientNetV2. EfficientNetV2 out-performed the other models by achieving a sensitivity of 0.77, specificity of 1.00, and F-measure of 0.77, demonstrating effectiveness in identifying true positives while minimizing false positives (Table 2).

**Table 2.** Comparative analysis of YouTube post classification using the proposed decision fusion module with adjustable confidence thresholds for species detection.

| **Metrics** | **Textual analysis** |  | **Visual Analysis** |  |  |
| --- | --- | --- | --- | --- | --- |
|  | **LSTM** | **All-MPNet-Base-V2** | **DarkNet53** | **InceptionResNetV2** | **EfficientNetV2** |
| Sensitivity | 0.45 | 0.69 | 0.66 | 0.68 | 0.77 |
| Specificity | 0.95 | 0.98 | 1.00 | 1.00 | 1.00 |
| Precision | 0.50 | 0.76 | 0.69 | 0.70 | 0.80 |
| Recall | 0.45 | 0.69 | 0.66 | 0.68 | 0.77 |
| F-measure | 0.47 | 0.72 | 0.66 | 0.67 | 0.77 |

The transformer model for text and EfficientNetV2 model for images were then used in conjunction to parallelly classify the text and image content of a post. The outputs were combined to assign three labels as (i) “Positive”-text model classified “Explicit Wildlife Trade” and image model detects presence of a species; (ii) “Uncertain” - when only one of the models gives a species/sales related output; and (iii) “Negative” when neither model gives a species/sales related output. Our findings revealed that the multimodal learning approach for wildlife trade detection exhibited a good degree of accuracy in identifying wildlife trade-related records (Figure 2). Notably, 77.42% of data points explicitly labeled as “Explicit Wildlife Trade” were correctly classified as “Positive,” underscoring the strong capability of the model in distinguishing relevant content. Furthermore, the proportion of instances where wildlife trade content was categorized as “Uncertain” (2.40%) or misclassified as “Negative” (3.85%) remains minimal, suggesting a low likelihood of erroneous classification. This outcome highlights the reliability of the approach in mitigating false negatives, a crucial factor in high-stakes applications such as illegal trade monitoring. Importantly, the model demonstrated high precision by ensuring that no data points categorized as “Irrelevant” content were mistakenly classified as “Positive.” This aspect is particularly significant, as it minimizes the risk of misclassification and enhances the overall trustworthiness of the system. Collectively, these results underscore the robustness and practical applicability of the proposed multimodal learning approach in accurately detecting and classifying wildlife trade content.

**Figure 2.**
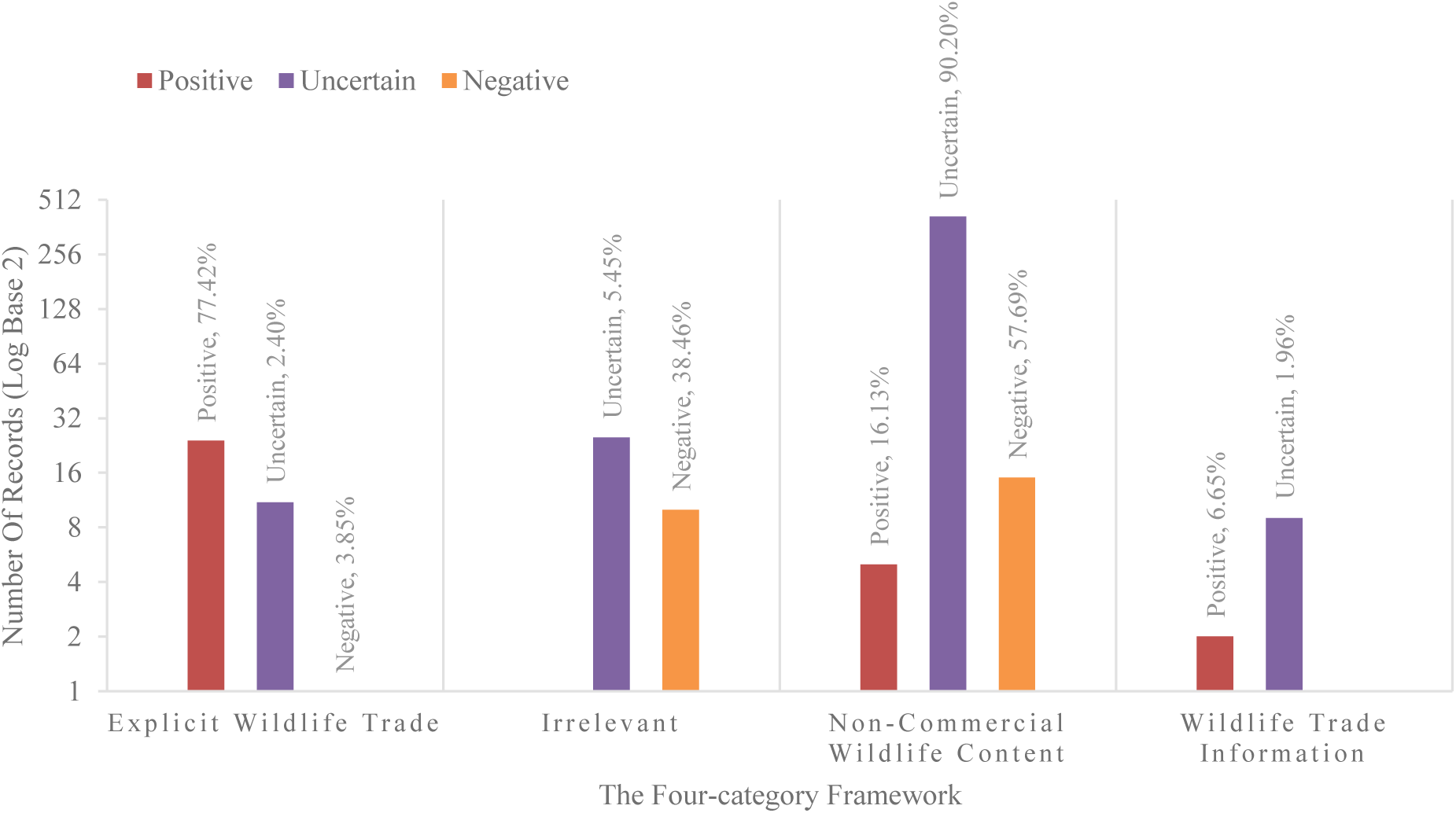
Evaluation of the multimodal learning approach for wildlife trade detection on test data.

Our analysis of 86,321 filtered YouTube posts identified 3,517 potential instances of wildlife trade involving 226 species, with a significant proportion belonging to the classes of birds, reptiles, and mammals (Figure 3). Notably, 51 species identified as “Explicit Wildlife Trade” are classified by the IUCN Red List as Critically Endangered. Within this group, birds account for the majority of the trade (722 instances), followed by reptiles (164 instances), and mammals (19 instances). Additionally, 62 species listed as Endangered were found in trade, encompassing 401 bird, 185 mammal, and 117 reptile trade instances, along with rare aquatic species such as 1 cartilaginous fish species and 47 ray-finned fishes. The category of Vulnerable species comprises 60 species, represented most heavily by birds (559 instances), again signaling the disproportionate targeting of avifauna across threat levels. Also notable are 81 mammals and 35 reptile trade instances under this category. A further 25 species are listed as Near Threatened, with 801 bird trade instances dominating the data. Finally, 28 species classified as Least Concern were also found in trade, including 418 bird instances.

**Figure 3.**
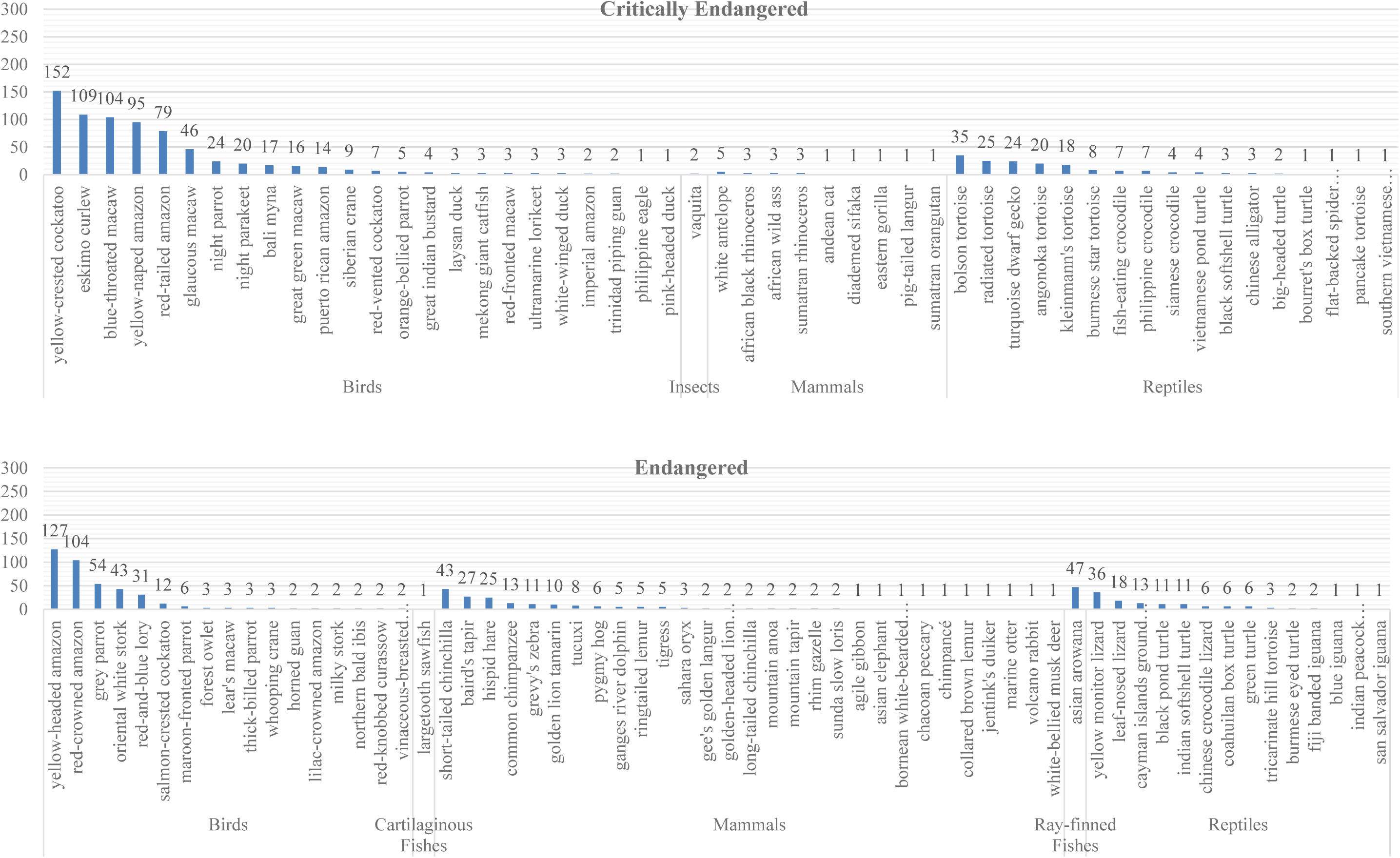
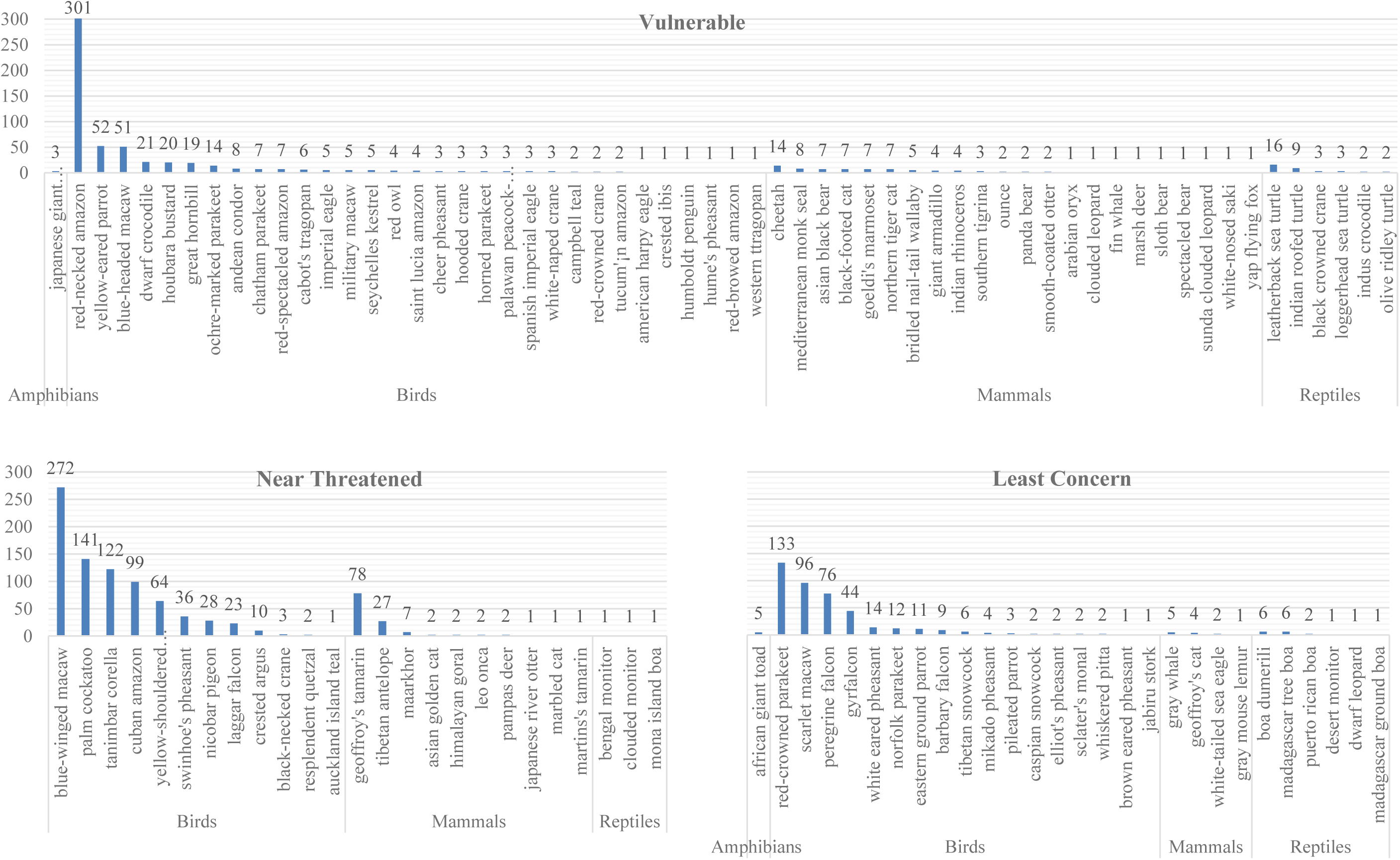
Results of the proposed multimodal learning approach for automated detection of species involved in the online wildlife trade. Species are classified according to the International Union for the Conservation of Nature (IUCN) Red List category they belong to, namely Least Concern, Near Threatened, Vulnerable, Endangered, Critically Endangered.

## Discussion

In this study, we introduce a method for video and textual content analysis of online wildlife trade. Our results demonstrate that this method more effectively identifies relevant content than unimodal text or video filtering because its decision fusion module synergistically combines visual and textual insights. This integration overcomes the inherent limitations of analyzing each modality independently. Similarly, previous work highlighted how online wildlife trade detection from images requires more than simple species identification (Kulkarni & Di Minin, 2023). Instances of trade in species at high risk of extinction were detected, highlighting that more research is now needed to investigate whether the trade in these species is legal or illegal and understand whether the individuals for sale were captive-bred or caught from the wild. The methods introduced here can also be used for other tasks in ecology and conservation science, such as tracking biodiversity trends and examining human-wildlife interactions reflected in social media.

One of the major challenges in detecting wildlife trade on platforms like YouTube is the sheer volume of data. By concentrating on certain keyframes and eliminating less pertinent ones, we substantially expedited the process of species identification within video content. The integration of both text and video filtering in our proposed method effectively identified and highlighted content related to wildlife trade. Furthermore, the integration of textual and visual analysis within a decision fusion module represents a significant advancement in our ability to detect wildlife trade. By implementing a decision fusion module to combine predictions from both modalities, our method ensures that only posts meeting specific criteria are classified as indicative of wildlife trade. This dual-layer validation is crucial, given the complexities inherent in detecting wildlife trade through nuanced textual phrases and visual representations.

Although our approach to keyframe analysis significantly reduces the computational burden associated with processing entire videos, it still requires substantial computational resources. The need to download and analyze video chunks, particularly for large datasets, can lead to time-consuming and resource-intensive processes. Optimizing these procedures and exploring alternative methods to reduce resource consumption remains a challenge. While fine-tuning pre-trained CNN models using the iNaturalist dataset improved accuracy, there remains a risk of overfitting to the specific characteristics of this dataset. The challenge lies in ensuring that the models generalize well across various contexts and species, particularly when exposed to new or unseen data. Continuous evaluation and adaptation of the models are necessary to maintain performance in dynamic real-world settings. Building upon this multimodal approach, future work may necessitate the implementation of a pipeline designed to operate over the complete dataset. This would enable a comprehensive and scalable analysis, ensuring that the benefits of this integrated methodology can be realized across a broader scope of information.

Wildlife trade is a multifaceted issue that encompasses legal and illegal activities, cultural practices, and varying definitions of what constitutes trade (‘t Sas-Rolfes et al., 2019). Capturing the nuances of this complexity through automated methods is inherently challenging. Posts may contain implicit references to trade or be situated within broader discussions of wildlife conservation, making it difficult for models to accurately classify such content without significant contextual understanding. The primary challenge in implementing a text filtering algorithm for wildlife trade detection lies in the ambiguity and variability of keywords. Keywords like “sale” and “dollar” can have multiple meanings, and the context in which they are used is crucial for accurate detection. Additionally, coded language, euphemisms, and regional differences can complicate the process (Davies et al., 2022). For this reason, future work should better consider these complicated aspects.

In conclusion, this study introduces one of the first examples of multimodal learning in conservation science. The same methods can be further enhanced by improving manual annotation of the data to include, for example, nuances such as legal versus illegal wildlife trade content. The proposed methods can also be used in iEcology to identify and count species from both photographs and text (Jarić et al., 2020). These simultaneous analyses of image and text caption may also be useful to assess the sentiment of social media posts than the image or text alone (Fink et al., 2020). Captions may provide detailed information on the species or activity, whereas the computer vision model may recognize only the broader class to which they belong.

## Methods

### Data collection

The multimodal learning approach begins by collecting various forms of metadata, such as post ID, channel name, channel link, title, description of the post, video ID, video link, and published time, from YouTube (Figure 1). A custom script was developed for automatically gathering and processing data from the YouTube Data Application Programming Interface (API) (Google, 2024) using common and scientific names of species listed in Appendix I of the Convention on International Trade in Endangered Species of Wild Fauna and Flora (CITES) (Nakamura & Kuemlangan, 2023). We chose Appendix I listed species because they are threatened by trade and international trade in these species is permitted only in exceptional circumstances and cannot be primarily for commercial use. The search terms were appended with trade-relevant keywords, such as “sale”, “purchase”, “dollar”, which are listed in Supplementary Appendix A. The library youtube_dl (version 2021.12.17) was used to download videos from YouTube for research purposes, allowing for customizable output paths and formats (youtube-dl, 2021). This resulted in the collection of a dataset comprising of 3,281,207 records, covering 523 mammal, cartilaginous fish, bony fish, bird, reptile, amphibian, and invertebrate species listed in Appendix I of CITES (see Supplementary Appendix B for a full list of species).

We defined a four-category framework to manually annotate a subset of the collected data. The “Explicit Wildlife Trade” class encompassed posts containing evidence of wildlife trade. This category included, for example, actual sales of wildlife. The “Wildlife Trade Information” class included posts that provided information or engaged in discussions about wildlife trade without directly promoting or participating in the activity. This category included, for example, news articles and educational content. The “ Other Wildlife Content “ class encompassed posts related to wildlife, but devoid of any trade implications, such as wildlife documentaries or conservation-focused discussions. The “Irrelevant” class was reserved for posts entirely unrelated to wildlife, for example a car for sale with a brand name of one of the species listed in Appendix I of CITES.

### Multimodal filtering

Following data collection, a multimodal filtering approach, combining textual and video filtering techniques, was used to filter out irrelevant posts. The textual feature extraction leveraged natural language processing (NLP) models to understand the meaning of the text content used in the post title and description, terms indicative of wildlife trade content. After filtering using text analysis, visual feature extraction was performed on remaining data which utilized deep convolutional neural networks (CNN) to detect visual patterns and elements in the video frames related to wildlife species. Finally, a subset of the data remaining after text and video filtering was used to construct a decision fusion module that combined the results from a separately trained NLP model (LSTM) and deep CNNs, enabling the automated identification of wildlife trade content on YouTube posts as a single system of classification. Random sub-samples of the data were manually annotated to verify and analyse each stage of the filtering method.

#### Text filtering

A text filtering algorithm was implemented that assigned a relevancy score between −1 and 1, to the text data for a particular post. A score of −1 indicating a probability of the post being highly irrelevant to wildlife trade, while 1 indicating the opposite. To calculate the relevancy score, we first developed two sets of template sentences which were based on sentences commonly found in the dataset. One set consisting of sentences representing the sale of species belonging to various taxa (positive template) and another set consisting of sentences representing the sale of non-animal entities (negative template). See Supplementary Appendix C for the list of templates. Sentences in the template were generated in two ways, (i) the authors crafted examples based on data that represented the general meaning of a post e.g. “animal for sale”/“car for sale”; and (ii) using a Large Language Model (LLM) based text generative algorithm from OpenAI, ChatGPT 3.5 (OpenAi). Using an LLM allowed us to generate varied example posts that mimicked the real data and increased the diversity of the template sets. This helped improve the efficiency of the method by increasing the pool of template sentences that carried the desired meaning of “animal for sale” vs “object for sale”. A similarity measure was calculated between the text from the data post and all the sentences from the template list. To calculate similarity, each sentence was first vectorised using a text vectorising algorithm ‘All-MPNet-Base-v2’ (Hugging-Face; Qureshi et al., 2021). Sentence vectorisation is a process that converts a sentence into a numerical array in a high dimensional geometric space that encodes the meaning of the contents of the sentence (Ul Haq et al., 2024). Thus, sentences that are similar to each other in meaning are found in the same vicinity within the geometric space. A numerical way to measure the similarity between two vectors is done by calculating the cosine of the angle between two vectors. Vectors that are identical have a cosine of 1 while those completely dissimilar have a cosine of 0. We then calculated the maximum cosine similarity between the data post and both the positive and negative templates. Finally, the relevancy score is calculated as:

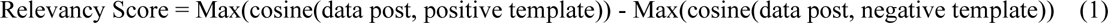

In principle, the Relavency Score is constructed in a way that if a data post is very similar to the positive template the relevancy score is greater than 0 and closer to 1, while if it is very similar to the negative template it is less than 0 and towards −1. Ambiguous posts, which measure a higher similarity to both the positive and negative templates, receive a score around 0. To finally mark a post as relevant or irrelevant, the relevancy score needed to be converted into a binary value (0/1). This was done using a subset of data that was manually annotated as “relevant” or “irrelevant”. The probability distributions of the relevancy scores were plotted for both the “relevant” and “irrelevant” subsets. As seen in Figure 4, the probability distributions of the two subsets are sufficiently separated, therefore validating the utility of the method. The point of intersection of the two distributions is selected as the threshold relevancy score. All data posts having a score below the threshold were designated as “irrelevant” and those equal or above were designated as “relevant”. Only data posts designated as “relevant” were passed on forward for further processing (Figure 1).

**Figure 4.**
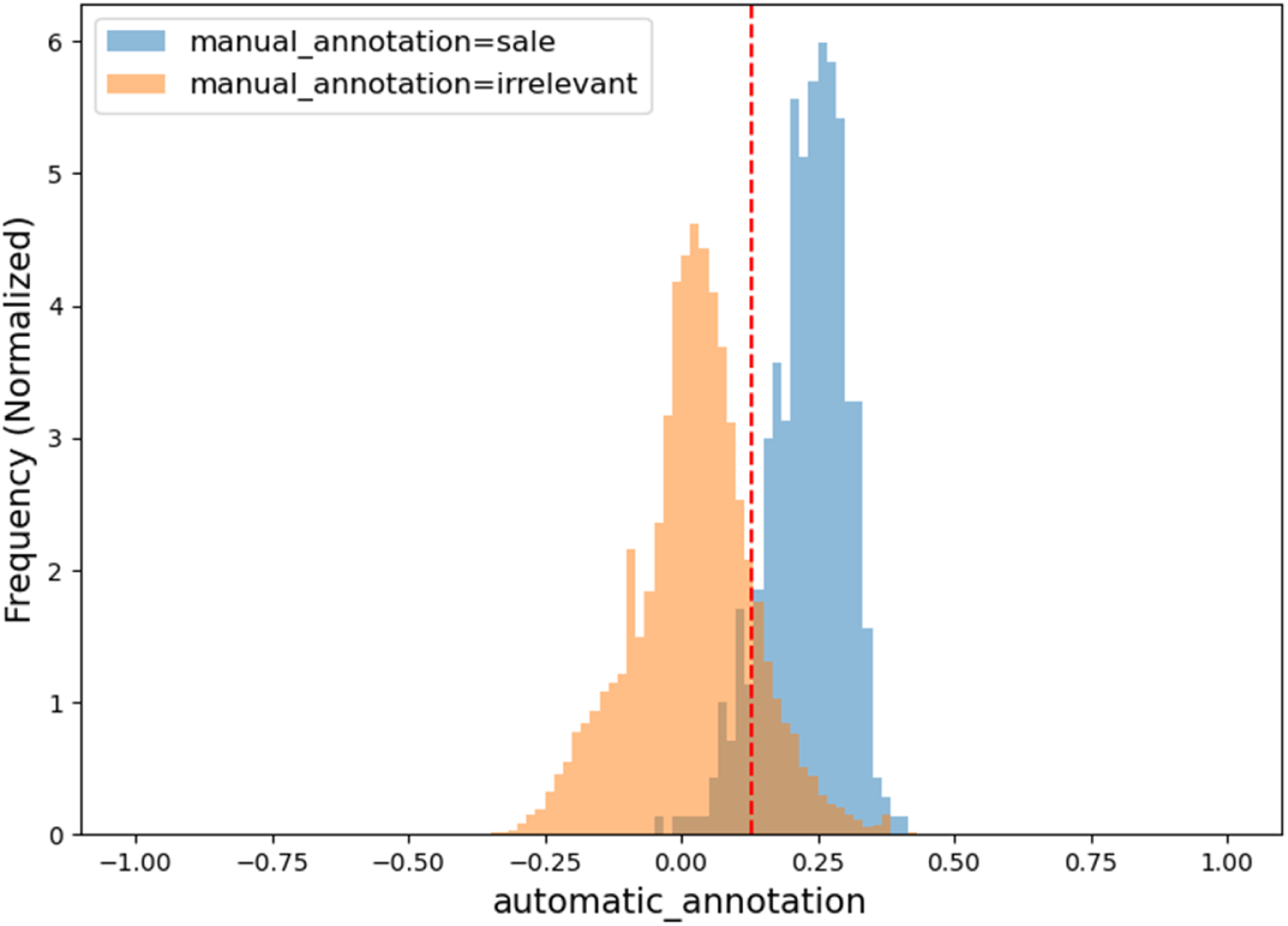
Probability distributions of relevancy scores for the “irrelevant” category compared to all other categories (“Explicit Wildlife Trade”, “Wildlife Trade Information”, and “Other Wildlife Content”) during the text filtering phase.

#### Video filtering

Following text filtering, the dataset was refined and forwarded to the video filtering stage, where records with unavailable videos were removed. However, analyzing all available video frames to extract wildlife-relevant information is resource-intensive, demanding substantial disk space, download time, and processing power due to their inherent size and redundancy. Thus, due to resource limitation, we selected 33,934 posts that had the highest score in text filtering. Video data is highly redundant due to several frames in a video being identical. This redundancy is particularly evident in the high degree of similarity between neighboring frames, a consequence of high frame rates and continuous motion present in video content (Ma et al., 2020). Therefore, we used a more computationally efficient approach that obviates the necessity to download and store the entire video file. Specifically, we prioritized the analysis of keyframes**—**representative frames that encapsulate the most salient visual content of a video (Tang et al., 2019)**—**rather than processing every individual frame. Furthermore, instead of downloading the entire video, we accessed it in smaller segments, or “chunks,” directly from the data stream. Keyframes were extracted from these chunks through a process involving three main steps: feature extraction, frame clustering, and keyframe selection. For feature extraction from downloaded frames, we evaluated pre-trained deep CNN models, namely DarkNet53 (Redmon & Farhadi, 2018), Inception-ResNet-v2 (Szegedy et al., 2017), and EfficientNetv2 (Tan & Le, 2021), and the optimal model was selected for further use. As the pre-trained models were not specifically designed for species classification, we enhanced their performance by fine-tuning them using the iNaturalist 2021 dataset (iNaturalist, 2021). This dataset encompasses millions of images across 10,000 species, providing taxonomic and geographic diversity that is crucial for robust image recognition models. Approximately 2.7 million images from this dataset were used for training and validation, with an additional test set of 100,000 images for evaluation. The models were trained using the Stochastic Gradient Descent (SGD) optimizer with a learning rate of 0.0001 and a momentum of 0.75. Training was conducted for 50 epochs. The model with the highest precision was selected for subsequent feature extraction. Fine-tuning the model with the iNaturalist dataset significantly enhanced its ability to detect the presence or absence of species. The extracted features represented high-level visual content of the frames.

We then clustered these features using the K-Means algorithm, grouping similar frames based on their visual similarity to identify redundant content or similar scenes. K-Means clustering has been heuristically applied with k values of 3, 4, and 5 to determine the optimal number of clusters. From each cluster, a representative frame was selected as the keyframe, summarizing the characteristics of that group. Subsequently, the fine-tuned CNN model assigned a prediction score to each keyframe indicating the probability of containing species. Finally, videos were filtered by applying a threshold to the species prediction scores of the video. A particular video was represented by its keyframes and the median of the prediction score over the 3 key frames was used as the representative score for that video. The threshold, above which videos were retained, was selected as the value of the confidence that minimised the number of “irrelevant” records in the manually annotated subset of the data. See Figure 5 for the distribution of median confidence scores for the “irrelevant” category as opposed to all other categories.

**Figure 5.**
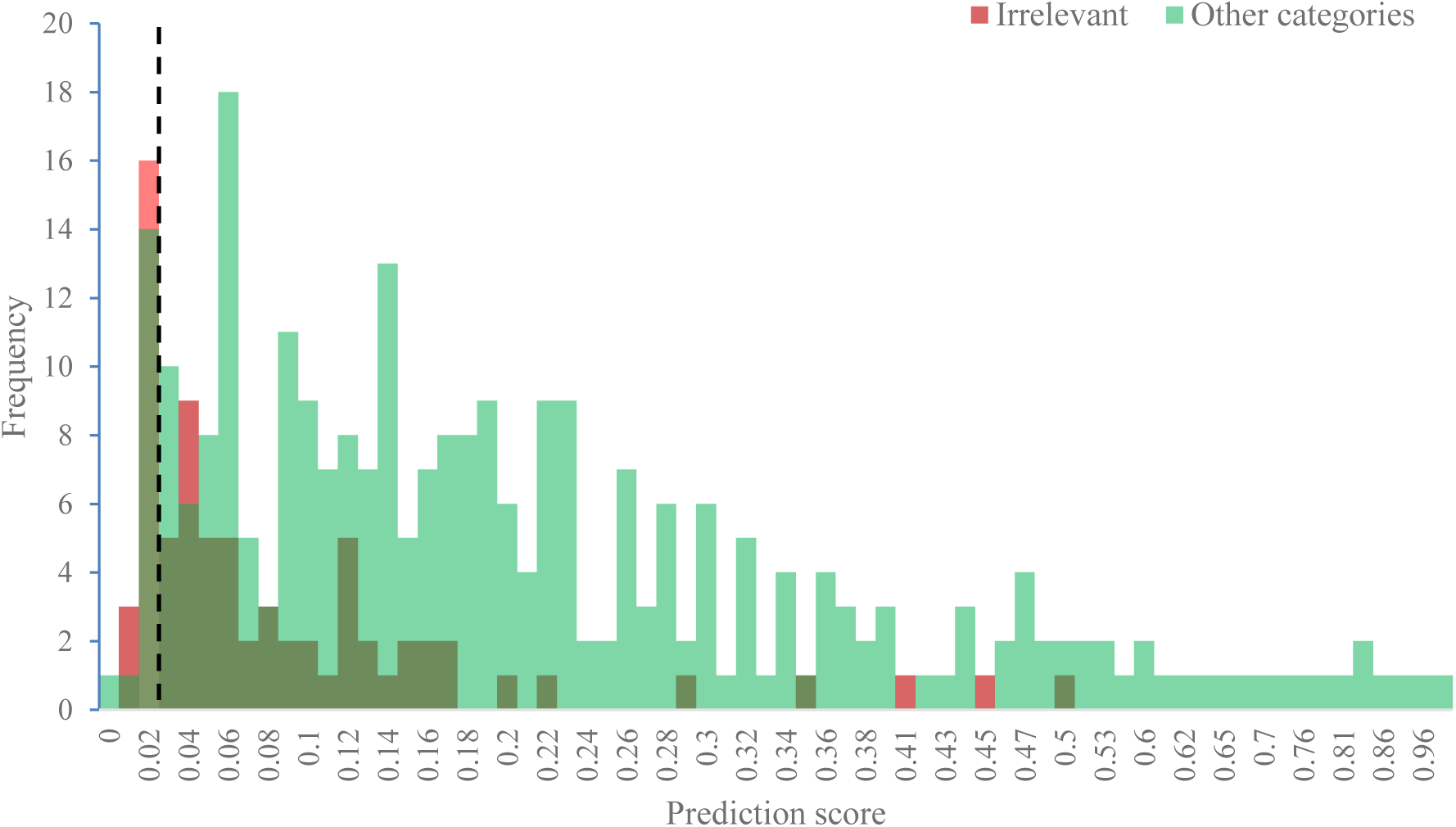
Probability distributions of median prediction scores for the “irrelevant” category compared to all other categories (“Explicit Wildlife Trade”, “Wildlife Trade Information”, and “Other Wildlife Content”) during the video filtering phase.

By identifying keyframes and filtering out videos based on a confidence threshold, our approach significantly reduced computational overhead, including disk input/output, network bandwidth, and CPU utilization, while effectively identifying videos containing relevant information. This method is particularly beneficial for large-scale video analysis tasks, where processing entire videos would be computationally infeasible.

#### Obtaining and assessing the filtered data

Evaluating the accuracy of the proposed multimodal filtering approach across the entire dataset is not only computationally intensive and time-consuming but also unfeasible due to the scale of the data. To address this, we sampled the data into 3 subsets, (i) before applying any filtering, (ii) after text filtering, and (iii) after video filtering. The random subsamples allowed us to get an estimate of the performance of the algorithms in filtering the desired categories. The size of the sample affects the precision and generalizability of the measures, underscoring the importance of careful sample size determination. We calculated the appropriate sample size using established methods for estimating population proportions based on margin of error and confidence intervals. Using the standard formula for sample size calculation (Garthwaite et al., 2024; Neyman, 1937):

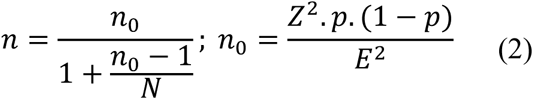

Given the absence of prior knowledge about the population proportion (*p*), it is standard practice to assume *p* = 0.5 (Altman, 2005). With the population size (*N*) of 3,281,207, a confidence level of 94%, a margin of error (*E*) of 5%, and Z-score (*Z*) of 1.9 corresponding to the desired confidence level, the calculated sample size (*n*) is approximately 353.

The text filtering method was applied to all 3,281,207 records, reducing the number to 495,533 using the optimal threshold relevance score of 0.13 (refer to Figure 4). However, unlike text data such as titles and descriptions, downloading full videos from YouTube presents significantly greater challenges due to the large data volumes and the bandwidth required. The YouTube Data API imposes strict limitations on video downloads, primarily to manage data usage and maintain platform stability. These limitations are enforced through rate limits and daily quotas, which restrict the amount of data that can be retrieved in a given period. To overcome these challenges, we designed the scope of our study to balance YouTube’s API restrictions with our need to gather a representative sample. By selecting 160,000 highest-scoring posts using text filtering, we were able to adhere to API’s constraints while obtaining a suitable number of videos for our research. The video filtering method was applied to 160,000 records, resulting in 89,642 videos after the removal of 24,235 unavailable videos and 21,888 duplicates. Subsequently, by applying the optimal threshold prediction score of 0.02 (refer to Figure 5), the number of videos was further reduced to 86,321.

### Multimodal classification

The filtered data was used to develop a single algorithm that can receive inputs both in the form of text and image to determine the relevancy of the post for wildlife trade. NLP models alone may not effectively identify which species are for sale based on text phrases, such as “the value of this is $979,” “the price is negotiable,” or “including special discount.” Likewise, using deep CNNs alone for wildlife trade detection can lead to inaccuracies, as keyframes might not explicitly indicate that species are for sale. Our objective was to integrate the outputs of both NLP models and deep CNNs to enhance the performance of wildlife trade detection. The workflow for textual and visual processing begins by initializing NLP and CNN models using PyTorch (version 1.10.0). To train and evaluate the models, we further manually annotated a subset of the filtered dataset, consisting of 3,437 records, using our four-category framework. These annotated records were then divided into three distinct subsets: 70% for training the model, 15% for validation purposes, and 15% for testing its performance. The outlined process for the proposed multimodal classification method is illustrated in Figure 1.

#### Classification of textual features

The textual feature extraction (Liang et al., 2017) is the process of transforming raw text data into numerical feature vectors suitable for subsequent text classification. In this study, we investigated two distinct models for feature extraction: the All-MPNet-Base-v2, a transformer-based model, and LSTM, a recurrent neural network model. Both models were trained on our four-category framework: “Explicit Wildlife Trade”, “Wildlife Trade Information”, “Other Wildlife Content”, and “Irrelevant”. To initiate feature extraction, both models first broke down the input text into smaller units called tokens. This process is called tokenization. The tokenizer used a word-piece tokenization approach, which splits words into subword units. This helps the models handle words that are not in their vocabulary. The vocabulary size is 30,527 tokens, and the tokenizer can process a maximum sequence length of 512 tokens. The tokenizer is case-sensitive, meaning it treats uppercase and lowercase characters differently.

The All-MPNet-Base-v2 model, a transformer-based architecture, then processed the tokenized text through multiple transformer layers. These layers are designed to analyze the relationships between tokens, capturing the contextual information within the text. The model comprises 12 layers, each with a hidden size of 768. The maximum tokenization length for the model is set to 512 tokens. The final output of the All-MPNet-Base-v2 model is a dense numerical vector, or embedding, that represents the text in a high-dimensional space. In this space, texts with similar semantic meanings are positioned closer together, facilitating accurate comparison and classification of text entries based on their content.

In contrast, the LSTM model, a recurrent neural network, processes the tokenized text sequentially. This sequential processing, combined with an internal memory state, allows the LSTM model to capture dependencies between words across the entire text sequence. The final hidden state of the LSTM network serves as the feature representation of the text. By employing these two distinct feature extraction models, we aimed to evaluate their effectiveness in capturing relevant features from text data for the task of wildlife trade detection. The model that demonstrated superior performance in accurately classifying text entries as relevant or irrelevant to wildlife trade was selected as the foundation for the final detection method. The training process for an LSTM-based text classifier uses a hidden layer dimension of 128 and two LSTM layers. During training, the best model is saved based on validation accuracy, and final evaluation is performed on the test set.

During the training phase, we employed oversampling and data augmentation techniques to balance the class distribution across the four-category framework. This involved identifying the most frequent class and randomly duplicating samples from the less represented classes. Linguistic transformations were then applied to these duplicated samples until the class distributions were balanced. These transformations included synonym replacement (substituting words with their synonyms), random insertion (inserting random synonyms at arbitrary positions), random swap (switching the positions of two words), and random deletion (removing words). By addressing class imbalance and exposing the model to a broader range of linguistic variations, this approach enhances the model’s ability to learn more robust and generalizable features (Haralabopoulos et al., 2021).

#### Classification of visual features

As detailed in the ‘video filtering’ section, we fine-tuned advanced deep CNN models (i.e., DarkNet53, Inception-ResNet-v2, and EfficientNetv2) using the iNaturalist 2021 dataset. To enhance the accuracy and generalization of these models, we employed various data augmentation techniques to expand and diversify the training dataset. These data augmentation techniques were integrated into the training pipeline using PyTorch’s *‘transforms.compose,’* which sequentially applied each transformation to the input images. Initially, images were resized to align with the input dimensions required by each model. We then implemented random cropping to focus the models on various regions of interest within the images. To account for variations in object orientation, horizontal flipping was applied, which involved creating mirrored versions of the images, allowing the models to become proficient in identifying objects presented in different orientations. Rotational transformations were also integrated into the process, where images were randomly rotated within a range of −30 to +30 degrees. Additionally, we employed affine transformations such as translation, scaling, and shearing to further enhance the robustness of the models. Translation involved shifting the images horizontally and vertically by up to 2% of their dimensions, helping the models to be resilient to positional variations. Scaling adjusted the size of the objects within the images within a range of 90% to 110%, simulating different distances from the camera, while shearing was applied with a maximum shear angle of 5 degrees, introducing slant distortions. To simulate varying lighting conditions encountered in natural environments, we incorporated color jittering. This technique involved randomly altering the brightness, contrast, saturation, and hue of the images, with variations set at ±10% for each attribute. Moreover, we introduced Gaussian noise to the images as part of the augmentation strategy. The Gaussian noise was applied with a mean of 0 and a standard deviation of 0.001, simulating the random variations in pixel intensities that often occur in real-world video data.

By incorporating these diverse and comprehensive data augmentation techniques into our training process, we aimed to build models that could learn robust and generalizable features. This approach was designed to make the models less sensitive to specific image characteristics, such as object orientation, lighting, and noise.

#### Integration of textual and visual analysis

The decision fusion module integrates the classifications from both textual and visual neural network outputs of YouTube posts to make a combined prediction based on learned features of both text and image. As previously mentioned, the classification of textural features involved employing NLP models to analyze the texused in the post, including the title and description. Species identification in visual analysis have been determined by applying an optimal threshold of 0.33 to the prediction score. This threshold was established based on the distribution of maximum prediction scores for the categories “Explicit Wildlife Trade,” “Wildlife Trade Information,” “Other Wildlife Content,” and “Irrelevant,” as illustrated in Figure 6. The Decision Fusion Module was designed using *if-then-else* logic to combine the outputs from two modalities. This approach enables the system to arrive at a final decision by applying specific criteria, ensuring the results of the textual and visual classifications are integrated for a coherent outcome. The decision fusion module assigns a “Positive” label to a YouTube post when both of the following conditions are satisfied: i) textual analysis detects wildlife trade, and ii) visual analysis identifies a species. If either or neither condition is met, the post is classified as “Uncertain” or “Negative,” respectively. “Negative” signifies high certainty that a post lacks wildlife trade content, assigned when neither textual nor visual analysis detects relevant information. Both modalities fail to meet their thresholds, allowing for confident exclusion from further analysis. “Uncertain” indicates partial evidence, applied when only one modality detects a relevant signal. This occurs when text suggests trade but visuals lack recognizable species, or when video shows potentially traded species without trade-related text. This intermediate label suggests the possibility of trade, warranting further review or manual validation due to the signal in one modality. To evaluate the models, six metrics were used including Sensitivity, Specificity, Precision, Recall, and F-Measure (Eq. 2 to Eq. 6, respectively).

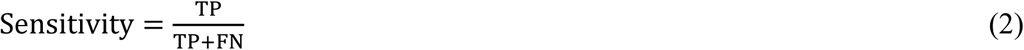

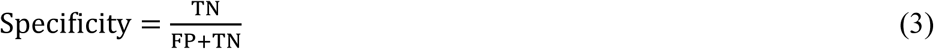

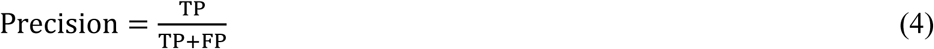

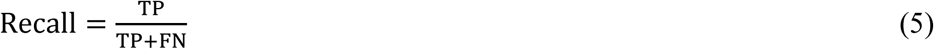

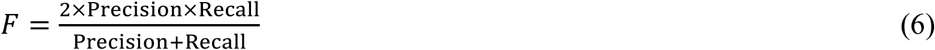

Where, TP, True positive; TN, True negative; FP, False positive; FN, False negative. The source code for reproducing the analyses and experiments presented in this study is available at: http://datadryad.org/share/XbOlrTatR89fDwbyH1elKGDnIJJq31qOAf6kvS7lI1M

**Figure 6.**
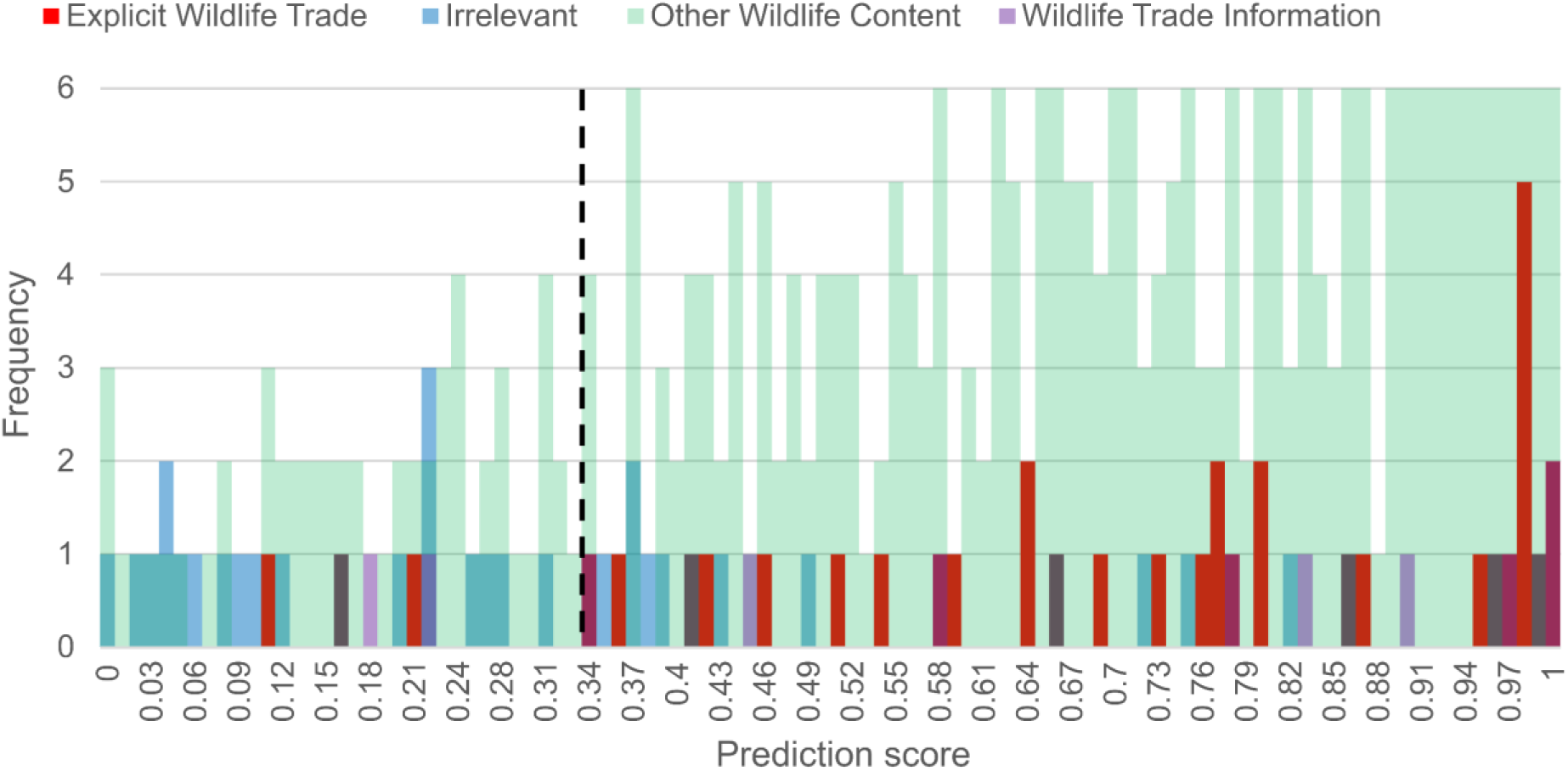
Probability distributions of maximum prediction scores for “Explicit Wildlife Trade,” “Wildlife Trade Information,” “Other Wildlife Content,” and “Irrelevant” subsets in the video filtering phase.

## Acknowledgements

E.D.M. was funded by the European Union (ERC, BIOBANG, 101171602). Views and opinions expressed are however those of the author(s) only and do not necessarily reflect those of the European Union or the European Research Council Executive Agency. Neither the European Union nor the granting authority can be held responsible for them. R.K. was funded by the KONE Foundation for this work (research grant 202103830). A.S.-R. was supported by a Ramón y Cajal fellowship (contract reference RYC2023-043755-I) from MICIU/AEI/10.13039/501100011033 and FSE+.

## Notes

### Competing Interest Statement

The authors have declared no competing interest.

## References

‘t Sas-Rolfes, M., Challender, D. W. S., Hinsley, A., Veríssimo, D., & Milner-Gulland, E. J. (2019). Illegal Wildlife Trade: Scale, Processes, and Governance. Annual Review of Environment and Resources, 44(Volume 44, 2019), 201–228. 10.1146/annurev-environ-101718-033253

Altman, D. G. (2005). Why We Need Confidence Intervals. World Journal of Surgery, 29(5), 554–556. 10.1007/s00268-005-7911-0

Cardoso, A. S., Bryukhova, S., Renna, F., Reino, L., Xu, C., Xiao, Z., Correia, R., Di Minin, E., Ribeiro, J., & Vaz, A. S. (2023). Detecting wildlife trafficking in images from online platforms: A test case using deep learning with pangolin images. Biological Conservation, 279, 109905. 10.1016/j.biocon.2023.109905

Cardoso, P., Amponsah-Mensah, K., Barreiros, J. P., Bouhuys, J., Cheung, H., Davies, A., Kumschick, S., Longhorn, S. J., Martínez-Muñoz, C. A., Morcatty, T. Q., Peters, G., Ripple, W. J., Rivera-Téllez, E., Stringham, O. C., Toomes, A., Tricorache, P., & Fukushima, C. S. (2021). Scientists’ warning to humanity on illegal or unsustainable wildlife trade. Biological Conservation, 263, 109341. 10.1016/j.biocon.2021.109341

Carvalho, A. F., de Morais, I. O. B., & Souza, T. B. (2023). Profiting from cruelty: Digital content creators abuse animals worldwide to incur profit. Biological Conservation, 287, 110321. 10.1016/j.biocon.2023.110321

Davies, A., D’Cruze, N., Senni, C., & Martin, R. O. (2022). Inferring patterns of wildlife trade through monitoring social media: Shifting dynamics of trade in wild-sourced African Grey parrots following major regulatory changes. Global Ecology and Conservation, 33, e01964. 10.1016/j.gecco.2021.e01964

Di Minin, E., Fink, C., Hiippala, T., & Tenkanen, H. (2019). A framework for investigating illegal wildlife trade on social media with machine learning. Conservation Biology, 33(1), 210–213. 10.1111/cobi.13104

Di Minin, E., Fink, C., Tenkanen, H., & Hiippala, T. (2018). Machine learning for tracking illegal wildlife trade on social media. Nature Ecology & Evolution, 2(3), 406–407. 10.1038/s41559-018-0466-x

Díaz, S., Settele, J., Brondízio, E. S., Ngo, H. T., Agard, J., Arneth, A., Balvanera, P., Brauman, K. A., Butchart, S. H. M., Chan, K. M. A., Garibaldi, L. A., Ichii, K., Liu, J., Subramanian, S. M., Midgley, G. F., Miloslavich, P., Molnár, Z., Obura, D., Pfaff, A., … Zayas, C. N. (2019). Pervasive human-driven decline of life on Earth points to the need for transformative change. Science, 366(6471), eaax3100. doi:10.1126/science.aax3100

Ek, C., & Samahita, M. (2023). Too much commitment? An online experiment with tempting YouTube content. Journal of Economic Behavior & Organization, 208, 21–38. 10.1016/j.jebo.2023.01.019

Fink, C., Hausmann, A., & Di Minin, E. (2020). Online sentiment towards iconic species. Biological Conservation, 241, 108289. 10.1016/j.biocon.2019.108289

Fink, C., Toivonen, T., Correia, R. A., & Di Minin, E. (2021). Mapping the online songbird trade in Indonesia. Applied Geography, 134, 102505. 10.1016/j.apgeog.2021.102505

Fukushima, C. S., Mammola, S., & Cardoso, P. (2020). Global wildlife trade permeates the Tree of Life. Biological Conservation, 247, 108503. 10.1016/j.biocon.2020.108503

Garthwaite, P. H., Moustafa, M. W., & Elfadaly, F. G. (2024). Locally correct confidence intervals for a binomial proportion: A new criteria for an interval estimator. Scandinavian Journal of Statistics, 51(1), 220–244. 10.1111/sjos.12672

Google, D. (2024). YouTube Data API. https://developers.google.com/youtube/v3

Haken, J. (2011). Transnational crime in the developing world. Global financial integrity, 32(2), 11–30.

Haq, R. U., Abdulabad, A., Asghar, S., & Szabo, J. K. (2023). Clicks and comments: Representation of wildlife crime in Pakistan in social media posts. Global Ecology and Conservation, 43, e02473. 10.1016/j.gecco.2023.e02473

Haralabopoulos, G., Torres, M. T., Anagnostopoulos, I., & McAuley, D. (2021). Text data augmentations: Permutation, antonyms and negation. Expert Systems with Applications, 177, 114769. 10.1016/j.eswa.2021.114769

Harrington, L. A., Auliya, M., Eckman, H., Harrington, A. P., Macdonald, D. W., & D’Cruze, N. (2021). Live wild animal exports to supply the exotic pet trade: A case study from Togo using publicly available social media data. Conservation Science and Practice, 3(7), e430. 10.1111/csp2.430

Hugging-Face. Sentence-transformers/all-mpnet-base-v2. Hugging Face, Accessed on 3 June 2024. https://huggingface.co/sentence-transformers/all-mpnet-base-v2

Hughes, L. J., Morton, O., Scheffers, B. R., & Edwards, D. P. (2023). The ecological drivers and consequences of wildlife trade. Biological Reviews, 98(3), 775–791. 10.1111/brv.12929

iNaturalist. (2021). iNaturalist community, https://www.inaturalist.org. Retrieved 29/04/2024 from

IUCN. (2001). IUCN Red List categories and criteria. International Union for the Conservation of Nature (IUCN).

Jarić, I., Correia, R. A., Brook, B. W., Buettel, J. C., Courchamp, F., Di Minin, E., Firth, J. A., Gaston, K. J., Jepson, P., Kalinkat, G., Ladle, R., Soriano-Redondo, A., Souza, A. T., & Roll, U. (2020). iEcology: Harnessing Large Online Resources to Generate Ecological Insights. Trends in Ecology & Evolution, 35(7), 630–639. 10.1016/j.tree.2020.03.003

Kulkarni, R., & Di Minin, E. (2021). Automated retrieval of information on threatened species from online sources using machine learning. Methods in Ecology and Evolution, 12(7), 1226–1239. 10.1111/2041-210X.13608

Kulkarni, R., & Di Minin, E. (2023). Towards automatic detection of wildlife trade using machine vision models. Biological Conservation, 279, 109924. 10.1016/j.biocon.2023.109924

Kulkarni, R., Wu, H., & Di Minin, E. (2025). Detection of trade in products derived from threatened species using machine learning and a smartphone. arXiv preprint, arXiv:2509.06585. 10.48550/arXiv.2509.06585

Lavorgna, A. (2014). Wildlife trafficking in the Internet age. Crime Science, 3(1), 5. 10.1186/s40163-014-0005-2

Liang, H., Sun, X., Sun, Y., & Gao, Y. (2017). Text feature extraction based on deep learning: a review. EURASIP Journal on Wireless Communications and Networking, 2017(1), 211. 10.1186/s13638-017-0993-1

Ma, S., Zhang, X., Jia, C., Zhao, Z., Wang, S., & Wang, S. (2020). Image and Video Compression With Neural Networks: A Review. IEEE Transactions on Circuits and Systems for Video Technology, 30(6), 1683–1698. 10.1109/TCSVT.2019.2910119

Marshall, B. M., Strine, C., & Hughes, A. C. (2020). Thousands of reptile species threatened by under-regulated global trade. Nature Communications, 11(1), 4738. 10.1038/s41467-020-18523-4

Moloney, G. K., Gossé, K. J., Gonedelé-Bi, S., Gaubert, P., & Chaber, A.-L. (2023). Is social media the new wet market? Social media platforms facilitate the online sale of bushmeat in West Africa. One Health, 16, 100503. 10.1016/j.onehlt.2023.100503

Morton, O., Scheffers, B. R., Haugaasen, T., & Edwards, D. P. (2021). Impacts of wildlife trade on terrestrial biodiversity. Nature Ecology & Evolution, 5(4), 540–548. 10.1038/s41559-021-01399-y

Nakamura, J. N., & Kuemlangan, B. (2023). Implementing the Convention on International Trade in Endangered Species of Wild Fauna and Flora (CITES) through national fisheries legal frameworks: A study and a guide–Second edition (Vol. 4). Food & Agriculture Org.

Neyman, J. (1937). Outline of a theory of statistical estimation based on the classical theory of probability. Philosophical Transactions of the Royal Society of London. Series A, Mathematical and Physical Sciences, 236(767), 333–380.

OpenAi. ChatGPT 3.5. In OpenAI. https://openai.com

Qureshi, A. H., Miao, Y., Simeonov, A., & Yip, M. C. (2021). Motion Planning Networks: Bridging the Gap Between Learning-Based and Classical Motion Planners. IEEE Transactions on Robotics, 37(1), 48–66. 10.1109/TRO.2020.3006716

Redmon, J., & Farhadi, A. (2018). Yolov3: An incremental improvement. arXiv preprint arXiv:1804.02767.

Rinne, J., Kulkarni, R., Soriano-Redondo, A., Correia, R., & Di Minin, E. Using automated content analysis to monitor global online trade in endemic reptile species. Diversity and Distributions, n/a(n/a). 10.1111/ddi.13771

Salas-Picazo, R. I., Ramírez-Bravo, O. E., Meza-Padilla, I., & Camargo-Rivera, E. E. (2023). The role of social media groups on illegal wildlife trade in four Mexican states: A year-long assessment. Global Ecology and Conservation, 45, e02539. 10.1016/j.gecco.2023.e02539

Scheffers, B. R., Oliveira, B. F., Lamb, I., & Edwards, D. P. (2019). Global wildlife trade across the tree of life. Science, 366(6461), 71–76. doi:10.1126/science.aav5327

Siriwat, P., & Nijman, V. (2018). Illegal pet trade on social media as an emerging impediment to the conservation of Asian otters species. Journal of Asia-Pacific Biodiversity, 11(4), 469–475. 10.1016/j.japb.2018.09.004

Soriano-Redondo, A., Alwasiti, H., Kulkarni, R., Correia, R. A., Bryukhova, S., Lita, N. M., Rigor, L. A., Tejerero, D. R., Tenazas, T. M., & Di Minin, E. (2023). Online wildlife trade in species of conservation concern. Conservation Letters, 16(6), e12985. 10.1111/conl.12985

Sung, Y.-H., Lee, W.-H., Leung, F. K.-W., & Fong, J. J. (2021). Prevalence of illegal turtle trade on social media and implications for wildlife trade monitoring. Biological Conservation, 261, 109245. 10.1016/j.biocon.2021.109245

Szegedy, C., Ioffe, S., Vanhoucke, V., & Alemi, A. (2017). Inception-v4, inception-resnet and the impact of residual connections on learning. Proceedings of the AAAI conference on artificial intelligence,

Tan, M., & Le, Q. (2021). Efficientnetv2: Smaller models and faster training. International conference on machine learning,

Tang, H., Liu, H., Xiao, W., & Sebe, N. (2019). Fast and robust dynamic hand gesture recognition via key frames extraction and feature fusion. Neurocomputing, 331, 424–433. 10.1016/j.neucom.2018.11.038

Tøn, A., Ahmed, A., Imran, A. S., Ullah, M., & Azad, R. M. A. (2024). Metadata augmented deep neural networks for wild animal classification. Ecological Informatics, 83, 102805. 10.1016/j.ecoinf.2024.102805

Ul Haq, I., Pifarré, M., & Fraca, E. (2024). Novelty Evaluation using Sentence Embedding Models in Open-ended Cocreative Problem-solving. International Journal of Artificial Intelligence in Education, 1–28.

Wang, Z., Chan, W.-P., Pham, N. T., Zeng, J., Pierce, N. E., Lohman, D. J., & Meng, W. (2023). One in five butterfly species sold online across borders. Biological Conservation, 283, 110092. 10.1016/j.biocon.2023.110092

Wyatt, T., Miralles, O., Massé, F., Lima, R., da Costa, T. V., & Giovanini, D. (2022). Wildlife trafficking via social media in Brazil. Biological Conservation, 265, 109420. 10.1016/j.biocon.2021.109420

Xu, Q., Cai, M., & Mackey, T. K. (2020). The illegal wildlife digital market: an analysis of Chinese wildlife marketing and sale on Facebook. Environmental Conservation, 47(3), 206–212. 10.1017/S0376892920000235

youtube-dl, c. (2021). youtube-dl. In (Version Version 2021.12.17) [Computer software]. GitHub. https://github.com/ytdl-org/youtube-dl

